# Polo-like kinase 1 prevents excess microtubule polymerization in *C. elegans* oocytes to ensure faithful meiosis

**DOI:** 10.1101/2024.08.03.606476

**Authors:** Juhi G. Narula, Sarah M. Wignall

## Abstract

Sexual reproduction relies on meiosis, a specialized cell division program that produces haploid gametes. Oocytes of most organisms lack centrosomes, and therefore chromosome segregation is mediated by acentrosomal spindles. Here, we explore the role of Polo-like kinase 1 (PLK-1) in *C. elegans* oocytes, revealing mechanisms that ensure the fidelity of this unique form of cell division. Previously, PLK-1 was shown to be required for nuclear envelope breakdown and chromosome segregation in oocytes. We now find that PLK-1 is also required for establishing and maintaining acentrosomal spindle organization and for preventing excess microtubule polymerization in these cells. Additionally, our studies revealed an unexpected new role for this essential kinase. While PLK-1 is known to be required for centrosome maturation during mitosis, we found that removal of this kinase from oocytes caused premature recruitment of pericentriolar material to the sperm-provided centrioles following fertilization. Thus, PLK-1 suppresses centrosome maturation during oocyte meiosis, which is opposite to its role in mitosis. Taken together, our work reveals multiple new roles for PLK-1 in oocytes, identifying PLK-1 as a key player that promotes faithful acentrosomal meiosis.

## INTRODUCTION

In most cell types, centrosomes mediate cell division by providing organizational cues for the formation of a microtubule-based bipolar spindle. Centrosomes are comprised of a centriole pair and surrounding pericentriolar material (PCM) that helps to both assemble the spindle by nucleating microtubules, and maintain bipolarity by organizing the microtubule minus ends into two poles. In contrast, oocyte meiosis is a specialized form of cell division in which centrosomes are typically absent. Nonetheless, bipolar spindles still form, demonstrating that alternate mechanisms must be utilized in oocytes to ensure faithful meiosis.

In the absence of centrosomes, mouse oocytes form multiple transient structures called acentriolar microtubule organizing centers (aMTOCs) that nucleate microtubules and facilitate bipolar spindle assembly (Schuh & Ellenberg, 2007). Interestingly, aMTOCs are not essential for spindle assembly, as spindles can still form in mouse oocytes depleted of these structures (So et al., 2022). Furthermore, aMTOCs have not been reported in human, bovine, or porcine oocyte spindles (So et al., 2022; Wu et al., 2022), further suggesting the dispensability of these structures in assembling a functional bipolar spindle. However, the mechanisms that guide acentrosomal spindle formation in the absence of aMTOCs remain unclear.

We utilized *C. elegans* oocytes as a model to reveal such mechanisms, since this system also does not rely on distinct MTOC structures for spindle formation (Connolly, Sugioka, Chuang, Lowry, & Bowerman, 2015; Wolff, Tran, Mullen, Villeneuve, & Wignall, 2016). Instead, microtubules nucleate in the vicinity of chromosomes upon nuclear envelope breakdown (NEBD) and form a cage-like structure. The microtubule minus ends are then sorted outwards to form multiple nascent poles that subsequently coalesce to form a bipolar spindle (Wolff et al., 2016). While this process is not centrosome-mediated, many factors contributing to proper spindle assembly during mitosis are utilized during meiosis. The kinase AIR-1^Aurora^ ^A^ facilitates PCM recruitment and centrosome maturation during mitosis (Hannak, Kirkham, Hyman, & Oegema, 2001) and is also required for spindle regulation during oocyte meiosis (Sumiyoshi, Fukata, Namai, & Sugimoto, 2015). Similarly, microtubule associated proteins TAC-1^TACC^ and ZYG-9^XMAP215^ regulate microtubule assembly at the centrosome (Bellanger & Gonczy, 2003; Le Bot, Tsai, Andrews, & Ahringer, 2003; Matthews, Carter, Thierry-Mieg, & Kemphues, 1998) and also stabilize the acentrosomal pole (Cavin-Meza et al., 2022; Harvey, Chuang, Sumiyoshi, & Bowerman, 2023). Thus, proteins found at the centrosome during mitosis can be repurposed during acentrosomal cell division to perform alternate functions.

Polo-like kinase 1 (PLK-1) is a highly conserved kinase that plays essential roles during mitosis in various organisms including mediating bipolar spindle assembly (Golsteyn, Mundt, Fry, & Nigg, 1995; Hartwell, Mortimer, Culotti, & Culotti, 1973; Llamazares et al., 1991; Sunkel & Glover, 1988) and PCM recruitment (Cabral, Laos, Dumont, & Dammermann, 2019; Ohta et al., 2021). Several targets of PLK-1 during mitosis have been identified, including the microtubule nucleating factor γ-tubulin and SPD-5, a scaffolding protein that helps build the PCM (Haren, Stearns, & Luders, 2009; Lane & Nigg, 1996; Rios et al., 2024; Woodruff et al., 2015). Additionally, PLK-1 mediates NEBD during *C. elegans* mitosis by phosphorylating nuclear lamina components (Chase et al., 2000; Martino et al., 2017; Rahman et al., 2015; Velez-Aguilera et al., 2020). While PLK-1’s roles during mitosis have been extensively studied, its functions in oocyte meiosis are less well characterized.

Work in mouse oocytes demonstrated that PLK1 is required for recruitment of centrosomal proteins to aMTOCs and for the subsequent assembly of a bipolar spindle (Clift & Schuh, 2015; Little & Jordan, 2020; Solc et al., 2015), thereby implicating PLK1 in aMTOC-driven spindle formation. A recent study of *C. elegans* oocytes reported spindle defects following PLK-1 inhibition, raising the possibility that this kinase contributes to aMTOC-independent spindle organization as well (Taylor et al., 2023). However, the specific roles that PLK-1 plays in spindle assembly and whether it is also necessary after spindle formation to ensure faithful meiosis remain unknown.

Here we address these questions by utilizing the auxin inducible degradation (AID) method to rapidly deplete PLK-1 from *C. elegans* oocytes. Using this system, we find that PLK-1 is required to maintain bipolar spindle stability, prevent excess microtubule nucleation throughout the oocyte, and inhibit premature maturation of the centrosome, thus revealing novel roles for PLK-1 in oocyte meiosis that are distinct from its functions in mitosis.

## RESULTS

### Auxin-mediated depletion of PLK-1 recapitulates known phenotypes

We first sought to test whether PLK-1 plays a role in assembling and stabilizing the acentrosomal spindle. A previous study demonstrated that PLK-1 depletion via RNAi inhibits nuclear envelope breakdown (NEBD) in oocytes thereby preventing spindle formation (Chase et al., 2000). Therefore, we utilized the auxin-inducible degradation (AID) system to achieve rapid protein depletion (Divekar, Horton, & Wignall, 2021; Zhang, Ward, Cheng, & Dernburg, 2015); we reasoned that this would allow us to deplete PLK-1 in a shorter time-frame after the nuclear envelope had already broken down in order to assess subsequent roles for PLK-1 during oocyte meiosis. Using CRISPR/Cas9, we generated a strain in which PLK-1 is tagged at the endogenous locus with both a GFP and degron tag on the N terminus, while the ubiquitin ligase TIR1 is expressed by a germline-specific promoter (**Figure 1A**). Notably, this strain exhibited low embryonic lethality (1.2% lethality; **Table 1**), indicating that the tags did not have a major effect on PLK-1 function and the TIR1 transgene did not negatively contribute to fitness. Additionally, we assessed the localization of the tagged PLK-1 protein and found that it localized to the centrosomes and holocentric kinetochores during mitosis, as well as to the chromosomes, spindle poles and midzone during various stages of oocyte meiosis (**Figure S1**), consistent with existing literature (Chase et al., 2000; Schmucker & Sumara, 2014; Taylor et al., 2023). Thus, the tags do not affect overall worm fitnesss or impair protein localization.

**Figure 1:**
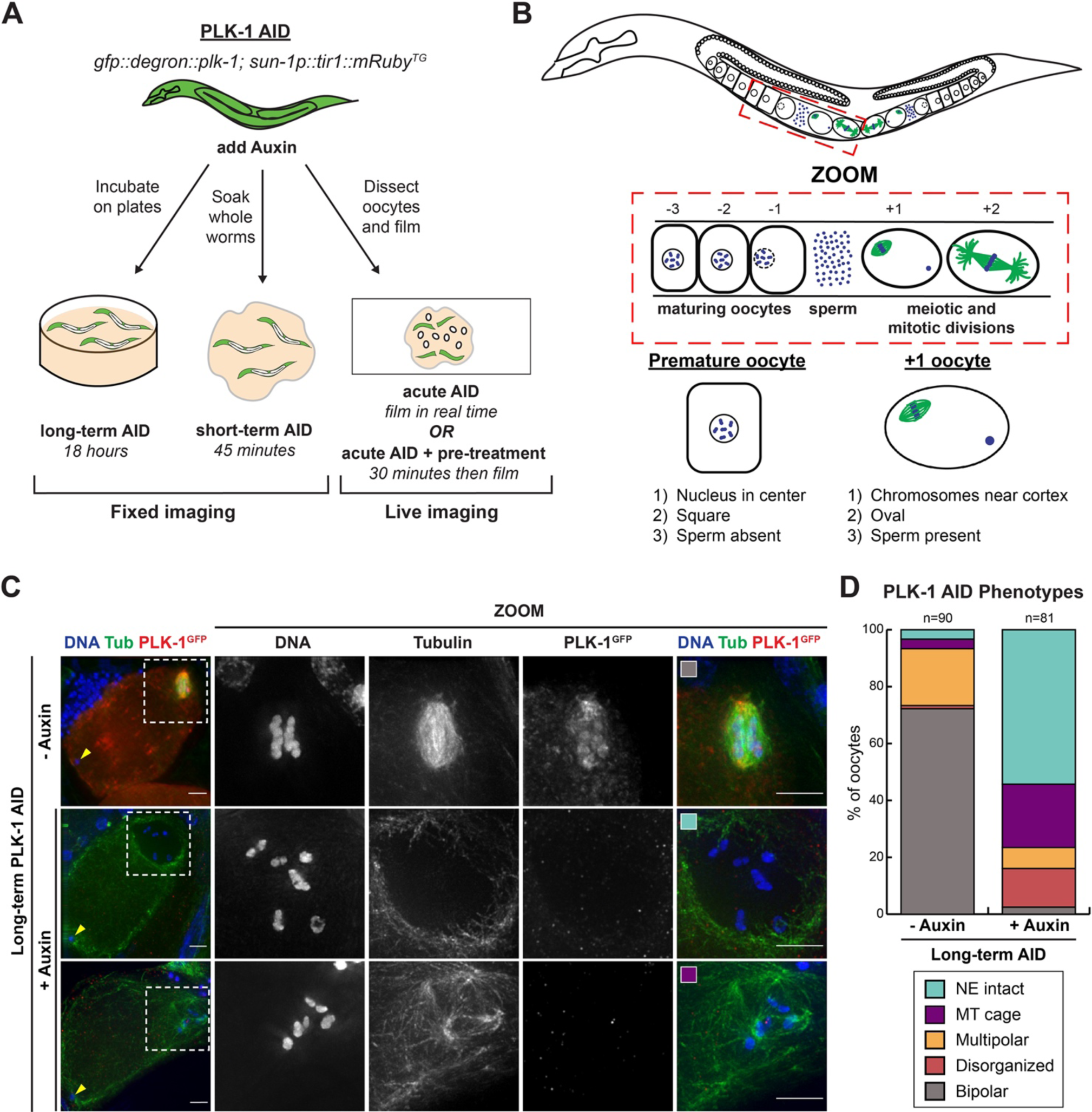
PLK-1 triggers nuclear envelope breakdown and the onset of meiosis. A) Schematic of the auxin-inducible degron (AID) strain that allows for germline-specific depletion of PLK-1 over various timescales in the presence of auxin. In this study, worms were incubated on auxin containing plates for 18 hours (long-term AID) or soaked in auxin for 45 minutes (short-term AID) and then fixed for immunofluorescence. Alternatively, worms were transferred into an auxin containing solution and oocytes were either immediately dissected (acute AID) or dissected after 30 minutes (acute AID + pre-treatment) and then imaged live. B) Oocytes in the *C. elegans* germ line mature as they progress towards the spermatheca, where they are fertilized and begin the meiotic divisions. To account for the loss of positional information during oocyte dissection we defined selection criteria to differentiate the +1 oocyte from premature oocytes, including the position of chromosomes, the shape of oocyte, and the presence of sperm. Only oocytes that were oval, fertilized, and had chromosomes near the cell cortex were included in the phenotype scoring shown in D. C) Control and long-term auxin-treated PLK-1 AID oocytes were stained for DNA (blue), tubulin (green) and PLK-1 (using a GFP antibody; red) and imaged at 40X magnification to view the entire cell, as well as 100X magnification to view the meiotic spindle (zooms). Arrowheads indicate sperm location as it corresponds to the selection criteria in 1B. Colored boxes correspond to phenotypic categories quantified in D. Scale bars = 5 μm. D) Quantification of the phenotypes observed in C. Without auxin, oocytes formed either multipolar or bipolar spindles, while long-term auxin treated oocytes either lacked tubulin density around the chromosomes (which we infer is caused by a NEBD defect) or form a microtubule (MT) cage around chromosomes, which reflects an early stage of spindle assembly. Oocytes with tubulin density around the chromosomes did not form properly organized bipolar spindles.

**Table 1:** Embryonic lethality & Brood size.

|  |  |
| --- | --- |
| **PHX3354** (*GFP::degron::plk-1*) control<br>n=10<br>Total eggs: 26<br>Total L1s: 2223<br>Embryonic lethality: 1.16%<br>Embryonic viability: 98.8% | **PHX3354** (*GFP::degron::plk-1*) + long-term AID<br>n=10<br>Total eggs: 947<br>Total L1s: 0<br>Embryonic lethality: 100%<br>Embryonic viability: 0% |

**Table 2:** *C. elegans* strains used:

| **Name** | **Description** | **Genotype** |
| --- | --- | --- |
| OC1002 | SPOT-tagged ZYG-1 strain | *zyg-1(bs197 [zyg-1::spot]) II* |
| PHX3354 | PLK-1 AID strain:<br>*GFP::degron::plk-1; Psun-1::TIR1::mRuby* | *unc-119(ed3) III; ieSi38 IV; plk-1(syb3354) [GFP::degron::plk-1] III* |
| PHX7793 | PLK-1 AID live imaging strain:<br>*GFP::degron::plk-1; Psun-1::TIR1::mRuby; mCherry::histone; GFP::tubulin* | *ItSi220[pOD1249/pSW077; Pmex-5::GFP::tbb-2-operon-linker-mCherry::his-11; cb-unc119(+)] I; unc-119(ed3), plk-1(syb7793) [GFP::degron::plk-1] III; ieSi38 [Psun-1::TIR1::mRuby::sun-1 3'UTR, Cbr-unc-119(+)] IV* |
| SMW44 | *mcherry::histone; GFP::tubulin; Psun-1::TIR1::mRuby* | *ItSi220[pOD1249/pSW077; Pmex-5::GFP::tbb-2-operon-linker-mCherry::his-11; cb-unc119(+)] I; unc-119(ed3) III; ieSi38 [Psun-1::TIR1::mRuby::sun-1 3'UTR + Cbr-unc-119(+)] IV.* |

To validate that we could recapitulate the known PLK-1 depletion phenotype using this strain, we incubated worms on plates containing 1 mM auxin for 18 hours (‘long-term AID’, **Figure 1A**). This treatment resulted in reduced PLK-1 staining and 100% embryonic lethality (**Figure 1C**, **Table 1**). We also observed a large number of oocytes with a hollow area of tubulin staining around the condensed chromosomes, consistent with the size of the nucleus (Wolff et al., 2016) (**Figure 1C**). Since previous work has shown that PLK-1 promotes nuclear envelope breakdown (Chase et al., 2000; Martino et al., 2017; Velez-Aguilera et al., 2020), we infer that this absence of tubulin density is due to the presence of an intact nuclear envelope. To ensure that these were not premature oocytes that had not yet triggered NEBD, we evaluated the oocytes using a number of criteria (**Figure 1B**). Maturing oocytes in the *C. elegans* germ line progress in an assembly-like fashion towards the spermatheca where they are fertilized. As oocytes mature the nuclear envelope breaks down and the meiotic spindle begins to form (McCarter, Bartlett, Dang, & Schedl, 1999). Since oocytes undergo morphological changes during this process, oocytes in the +1 position can be identified based on the position of the chromosomes relative to the cortex, the shape of the oocyte, and the absence of sperm (**Figure 1B**). We found that following long-term PLK-1 AID the majority of oocytes lacked a bipolar spindle, and instead displayed the empty region of tubulin staining (‘NE intact’; 44/81 oocytes, 54%), or a microtubule array where some tubulin density is present but no bipolar spindle has assembled (‘MT cage’; 18/81 oocytes, 22%) (**Figure 1D**). Our findings thus support previously published work implicating PLK-1 in promoting nuclear envelope breakdown, and validate the use of our AID strain to investigate PLK-1 function.

### PLK-1 is required for bipolar spindle assembly and stability

In our long-term AID experiments, we observed a subset of PLK-1-depleted oocytes where NEBD had occurred (19/81 oocytes; 23.5%) (**Figure 1D**). Notably, spindles in these oocytes were often characterized by excess poles or complete disorganization (“multipolar” and “disorganized” categories, respectively), suggesting that PLK-1 may be required for normal spindle assembly. This is in line with a recent study that reported spindle defects using a different method of PLK-1 inhibition (Taylor et al., 2023).

To better understand how PLK-1 might promote bipolar spindle assembly, we next attempted to remove PLK-1 from oocytes after NEBD and assess its effects on spindle organization. Soaking worms in a 5 mM auxin solution for 45 minutes (‘short-term AID’, **Figure 1A**) was sufficient to achieve a consistent loss of PLK-1 staining, and spindles that were able to form in these conditions exhibited various defects (**Figure S2A**), thus implicating PLK-1 in spindle assembly. However, this short-term AID treatment also resulted in a large percentage of oocytes with intact nuclear envelopes (42/68 oocytes; 61.8%, **Figure S2B**), similar to long-term AID (**Figure 1B**), suggesting that PLK-1 was being depleted prior to NEBD in many oocytes. Because we were unable to achieve robust PLK-1 depletion with shorter soaking times, we instead used RNAi to enrich for oocytes in which the nuclear envelope had already broken down by depleting the anaphase promoting complex component EMB-30. This condition arrests oocytes in metaphase I, allowing us to enrich for oocytes with assembled bipolar spindles (Furuta et al., 2000). Under these conditions, we found that the majority of auxin-treated spindles were either multipolar (13/89; 14.6%) or disorganized (61/89; 68.5%, **Figure 2A, 2B**), supporting a role for PLK-1 in promoting proper spindle architecture.

**Figure 2:**
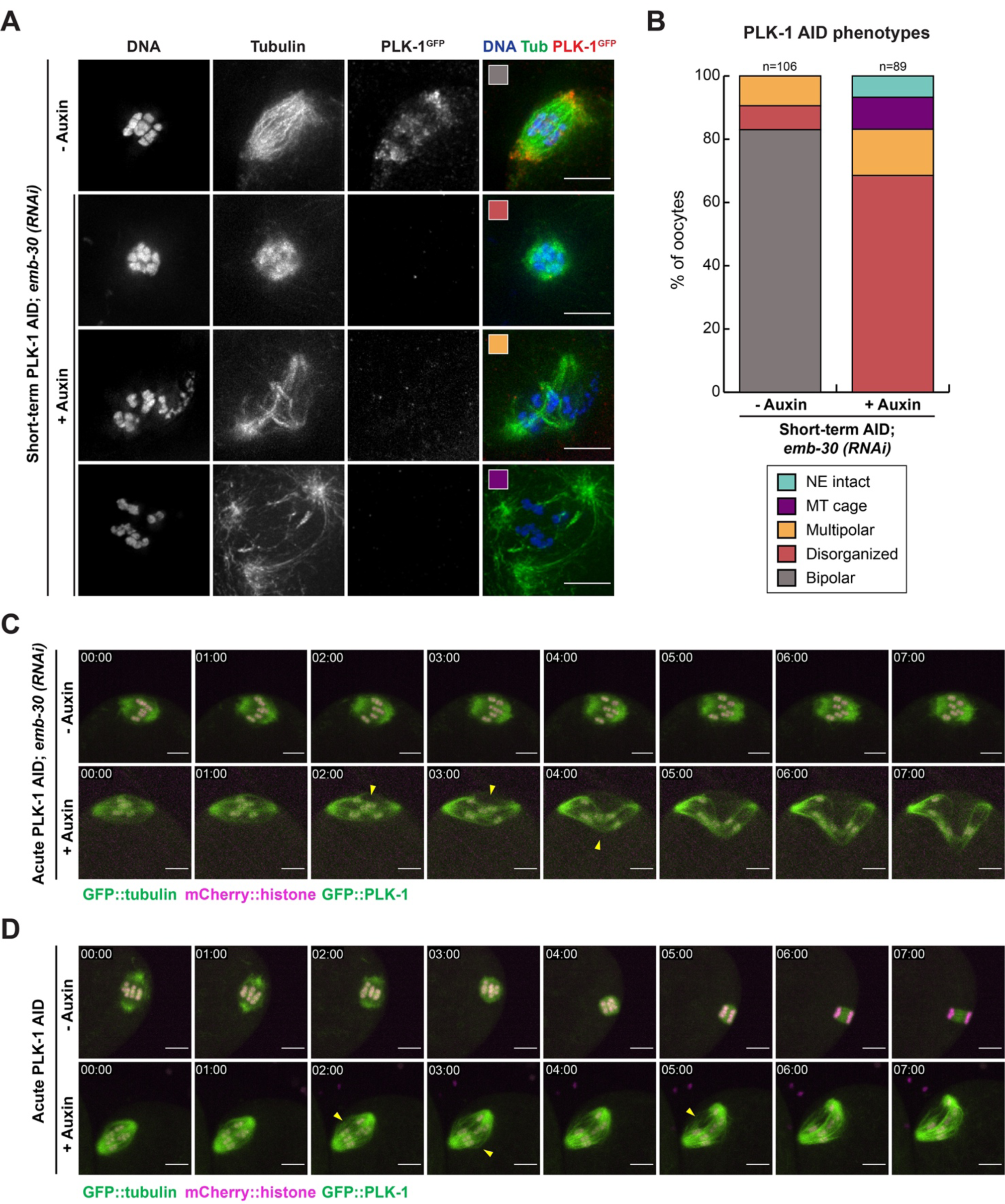
PLK-1 is required to maintain bipolar spindle stability. A) Control and short-term auxin treated PLK-1 AID *emb-30(RNAi)* metaphase-arrested oocytes were stained for DNA (blue), tubulin (green) and PLK-1 (using a GFP antibody; red). Representative images of the major phenotypic categories quantified in B are indicated with colored boxes. Scale bars = 5 μm. B) Quantification of the phenotypes observed in A. Control spindles were largely bipolar, while most short-term auxin treated oocytes were either unable to retain organization of the entire structure (disorganized) or of spindle poles (multipolar). Scale bars = 5 μm. C) *Ex utero* live imaging of *emb-30(RNAi)* metaphase-arrested PLK-1 AID oocytes expressing GFP::tubulin and GFP::PLK-1 (green) and mCherry::histone (magenta) after acute PLK-1 depletion. In control movies, the chromosomes oscillate and the spindle retains its integrity. In contrast, auxin treated spindles rapidly become disorganized, with poles moving apart and the spindle center losing integrity (arrowheads). Time elapsed shown in min:sec. Scale bars = 5 μm. D) *Ex utero* live imaging of unarrested oocytes expressing GFP::tubulin and GFP::PLK-1 (green) and mCherry::histone (magenta) after acute PLK-1 depletion. Control oocytes progress from metaphase where chromosomes are aligned, to anaphase where the chromosomes segregate bidirectionally. Upon acute AID the microtubules in the spindle center begin to splay (arrowheads) and the oocyte does not progress to anaphase. Time elapsed shown in min:sec. Scale bars = 5 μm.

Given that we are using metaphase arrest in these experiments, we infer that PLK-1 was depleted from many oocytes after the spindle had already achieved bipolarity, suggesting that PLK-1 is required to maintain spindle stability. To directly test this hypothesis, we generated a PLK-1 AID strain that also expressed mCherry::histone and GFP::tubulin to visualize the effects of PLK-1 removal from pre-formed spindles in real time. We performed *ex utero* live imaging of PLK-1 depletion by dissecting oocytes into a 500 µM auxin solution and immediately filming the spindles (‘acute AID’, **Figure 1A**). Live imaging of metaphase-arrested (**Figure 2C, Videos 1-2**) and unarrested oocytes (**Figure 2D, Videos 3-4**) revealed that spindles began to exhibit defects soon after PLK-1 depletion was initiated. In control conditions, metaphase-arrested spindles (7/7 spindles) as well as unarrested spindles (8/8 spindles) retained their structure. In contrast, when oocytes were dissected into auxin, the spindle poles moved apart while remaining relatively focused, and the midspindle region lost its integrity. This resulted in a disorganized structure during metaphase arrest (6/6 spindles) that in unarrested conditions was unable to progress to anaphase (7/7 spindles). Because we observed similar spindle defects in both arrested and unarrested oocytes, these phenotypes are likely not a byproduct of the metaphase arrest. Altogether, our results support a role for PLK-1 in both assembling and stabilizing the oocyte spindle.

### PLK-1 prevents the formation of ectopic microtubule asters throughout the oocyte

In characterizing the aberrant spindles that formed following PLK-1 AID, we assessed the localization of the microtubule minus end marker ASPM-1, which typically marks the two poles in control spindles (**Figure 3A**). This analysis demonstrated that some spindles were completely disorganized, with ASPM-1 present in a diffuse haze, while others had excess ASPM-1-marked spindle poles. Unexpectedly, we also found a novel phenotype not previously reported using other depletion methods (Chase et al., 2000; Taylor et al., 2023). Control treated oocytes typically do not have distinct concentrations of tubulin density or ASPM-1 in the cell other than at the meiotic spindle. However, after short-term (**Figure 3A, 3C**) and long-term (**Figure 3B, 3C**) PLK-1 depletion, we observed ectopic microtubule asters near the oocyte chromosomes that were marked by distinct populations of ASPM-1.

**Figure 3:**
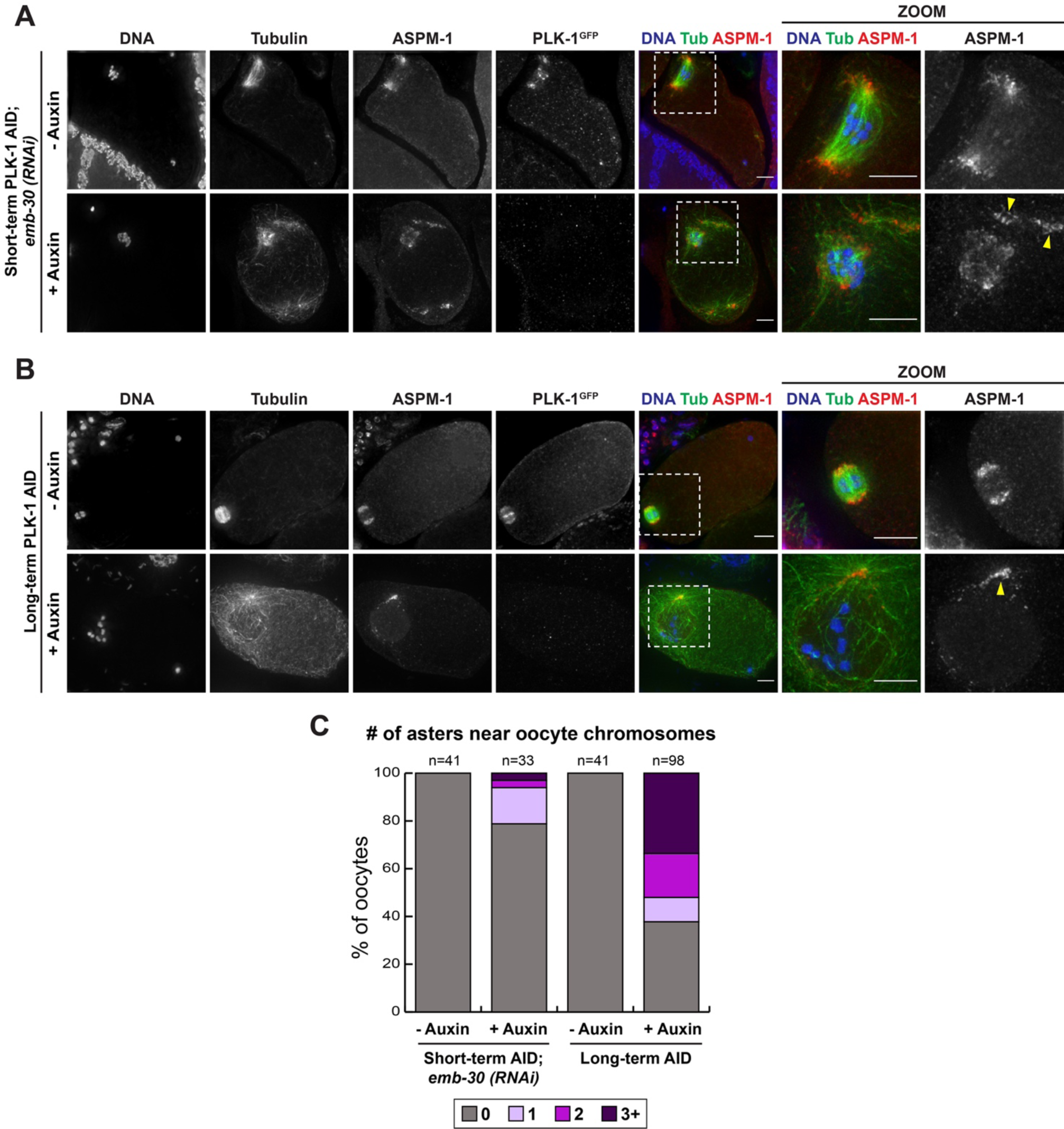
PLK-1 depletion results in disorganized spindles and excess microtubule polymerization near the oocyte chromosomes. A) Control and short-term auxin treated PLK-1 AID *emb-30(RNAi)* metaphase-arrested oocytes were stained for DNA (blue), tubulin (green), ASPM-1 (red) and PLK-1 (using a GFP antibody; not shown in merge) and imaged at 40X (columns 1-5) and 100X magnification (zooms). ASPM-1-positive microtubule asters can be observed in auxin-treated oocytes (arrowheads) and the minus ends of microtubules within the spindle are also disorganized, as evidenced by a haze of ASPM-1. Scale bars = 5 μm. B) Control and long-term auxin treated PLK-1 AID oocytes were stained for DNA (blue), tubulin (green), ASPM-1 (red) and PLK-1 (using a GFP antibody; not shown in merge) and imaged at 40X (columns 1-5) and 100X magnification (zooms). Microtubule density co-incident with ASPM-1 staining forms near the intact nuclear envelope (arrowhead). Scale bars = 5 μm. C) Quantification of the phenotypes observed in A and B. Both modes of PLK-1 depletion resulted in increased microtubule aster formation.

Curiously, we did not observe the formation of these microtubule asters following acute PLK-1 AID (**Figure 2C, 2D**, **4B** top row) when oocytes were dissected directly into auxin and then filmed for 15 minutes to assess immediate effects on the spindle. However, at the end of that timeframe, residual GFP::PLK-1 signal was often still apparent on the chromosomes (**Figure 2C**), indicating that we were not achieving complete depletion of PLK-1. This is consistent with the fact that we needed to soak worms in auxin for 45 minutes in our short-term AID experiments to observe complete loss of PLK-1 staining (**Figure 2A**). Thus, we modified our acute AID method to achieve more robust depletion in line with the short-term AID treatment. To this end, we soaked the worms in auxin for 30 minutes prior to dissection and then performed *ex utero* live imaging for 15 minutes (‘acute AID + pre-treatment’, **Figure 1A**). In control conditions, no significant tubulin density or asters were present throughout the oocyte (7/7 oocytes, **Figure 4A, Video 5**). Conversely, following auxin treatment ectopic asters formed throughout the oocyte and converged with each other over time, resulting in larger more tubulin-rich structures (7/7 oocytes, **Figure 4A, Video 6**). The lack of ectopic asters present after immediate administration of auxin suggests that this phenotype only arises after robust PLK-1 depletion, as compared to the spindle defects that were apparent even when PLK-1 was partially depleted from the oocyte (**Figure 2D**).

**Figure 4:**
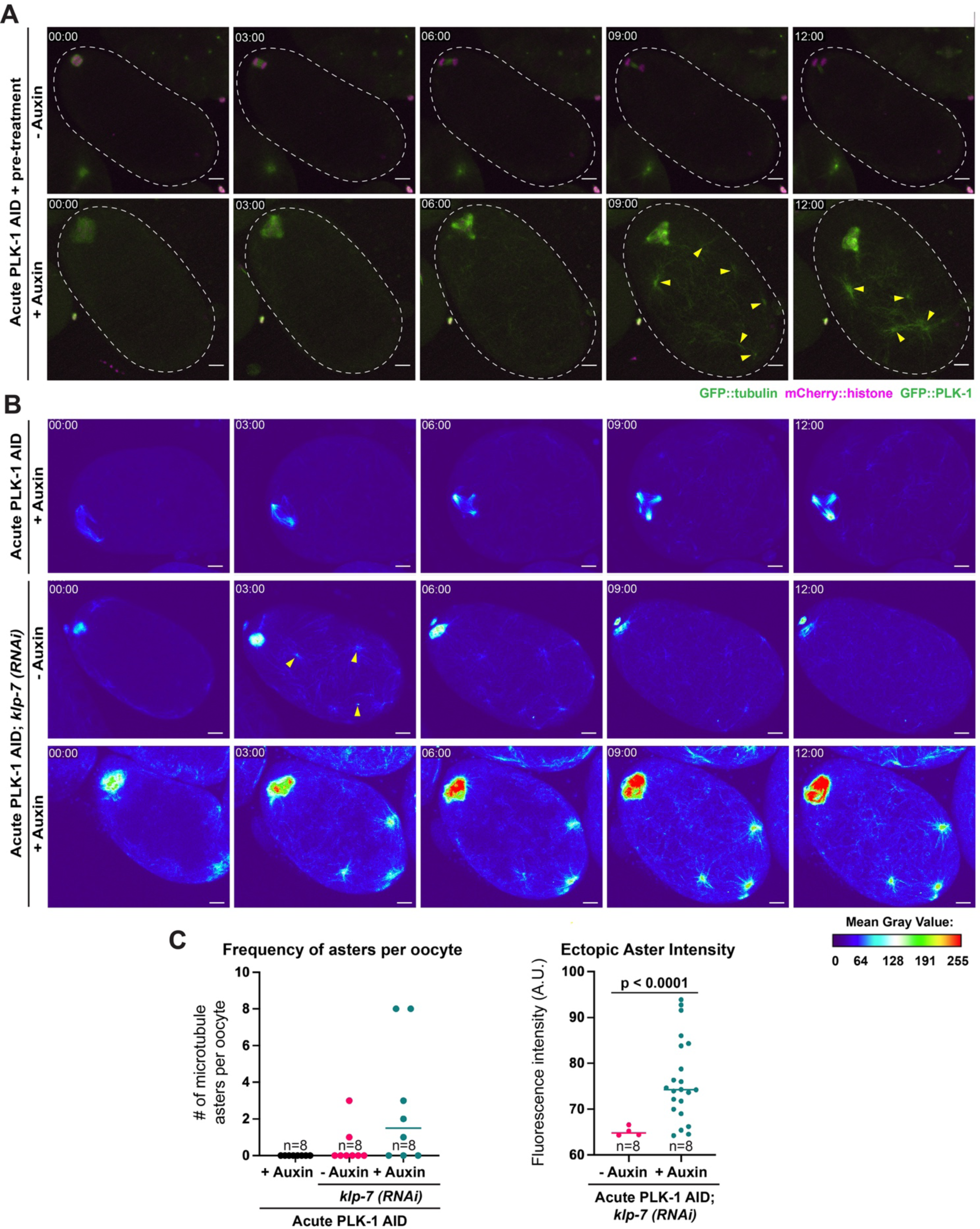
Dual depletion of PLK-1 and the microtubule depolymerase KLP-7 results in increased microtubule aster formation. A) *Ex utero* live imaging of entire oocytes (white dashed line) expressing GFP::tubulin and GFP::PLK-1 (green) and mCherry::histone (magenta) after acute AID + pre-treatment. Control oocytes (top row) were able to segregate chromosomes and complete meiosis I, and did not contain areas of strong tubulin density other than at the meiotic spindle. In comparison, PLK-1 depleted oocytes (bottom row) exhibited spindle defects and formed ectopic microtubule asters throughout the cell (arrowheads). Time elapsed shown in min:sec. Scale bars = 5 μm. B) *Ex utero* live imaging of PLK-1 AID oocytes expressing GFP::tubulin and GFP::PLK-1; images are pseudocolored to show the mean gray value of GFP intensity. Acute PLK-1 depletion alone resulted in no ectopic microtubule asters throughout the oocyte (top row). Following *klp-7(RNAi)*, oocytes had some instances of ectopic aster formation (arrowheads, second row), but microtubule density dramatically increased upon dual depletion of KLP-7 and PLK-1 (arrowheads, bottom row). Time elapsed shown in min:sec. Scale bars = 5 μm. C) Quantification of ectopic microtubule aster intensities in oocytes after 12 minutes of acute AID treatment. 8 oocytes were imaged in all conditions. The graph on the left shows the number of asters per oocyte that were above a threshold of 64 A.U.; the graph on the right shows the intensities of these asters. More asters were present in oocytes depleted of both KLP-7 and PLK-1 than in either single depletion, and these asters had increased fluorescence intensity compared to those occurring from KLP-7 depletion alone. Statistical significance was determined using a two-tailed unpaired t-test with Welch’s correction (t=5.599). **** = P<0.0001.

Previous work has shown that depletion of the microtubule depolymerase KLP-7^MCAK^ via RNAi results in the formation of ectopic microtubule asters (Gigant et al., 2017), reminiscent of the PLK-1 depletion phenotype. Moreover, there is evidence that PLK-1 phosphorylates KLP-7 during mitosis to promote its depolymerase activity (Zhang et al., 2011). Therefore, we hypothesized that the ectopic asters seen upon PLK-1 depletion might be caused by misregulation of KLP-7. If this were the sole mechanism by which PLK-1 AID ectopic aster formation was occurring, we would expect co-depletion of PLK-1 and KLP-7 to exhibit a similar phenotype as either single depletion alone. As expected, KLP-7 RNAi resulted in the formation of some ectopic microtubule asters (8/8 oocytes, **Video 7**) while acute PLK-1 AID alone did not result in ectopic aster formation (**Figure 4B, Video 8**). Strikingly, when we combined KLP-7 RNAi with acute PLK-1 AID, there was a significant increase in microtubule polymerization compared to either individual depletion condition alone (8/8 oocytes, **Figure 4B, 4C, Video 9**). These co-depleted oocytes had an increased number of ectopic microtubule asters, and the asters that formed had increased tubulin intensity compared to those that formed following *klp-7(RNAi)* alone (**Figure 4C**). Although these results do not rule out the possibility that PLK-1 regulates KLP-7 in oocytes, they suggest that the microtubule asters formed following PLK-1 AID are not solely caused by misregulation of this microtubule depolymerase.

### PLK-1 prevents premature recruitment of PCM to the sperm-provided centrioles

Interestingly, when analyzing microtubule polymerization in the short term and long-term PLK-1 depletion conditions, we often noticed excess microtubule density adjacent to the sperm DNA (**Figure 5A-C**). These asters were marked by distinct clusters of ASPM-1, suggesting that the minus ends are organized into distinct foci at these sites (**Figure 5A, 5B**). This ectopic microtubule polymerization was striking given what is known about the *C. elegans* germ line. During oogenesis, the maternal centrioles are eliminated prior to the turn of the gonad arm (Mikeladze-Dvali et al., 2012). Upon fertilization, the oocyte inherits a centriole pair from the sperm, but the sperm-provided centrioles do not recruit maternal PCM or nucleate microtubules until the completion of the meiotic divisions (McNally et al., 2012). This led us to speculate that the ectopic polymerization we observed near the sperm may be caused by premature centrosome maturation, implicating PLK-1 in regulating this process. Consistent with this hypothesis, we found that PLK-1 localizes to the sperm centrioles as demonstrated by its colocalization with ZYG-1, an essential kinase regulating centriole replication (O’Connell, 2002) (**Figure 5D**).

**Figure 5:**
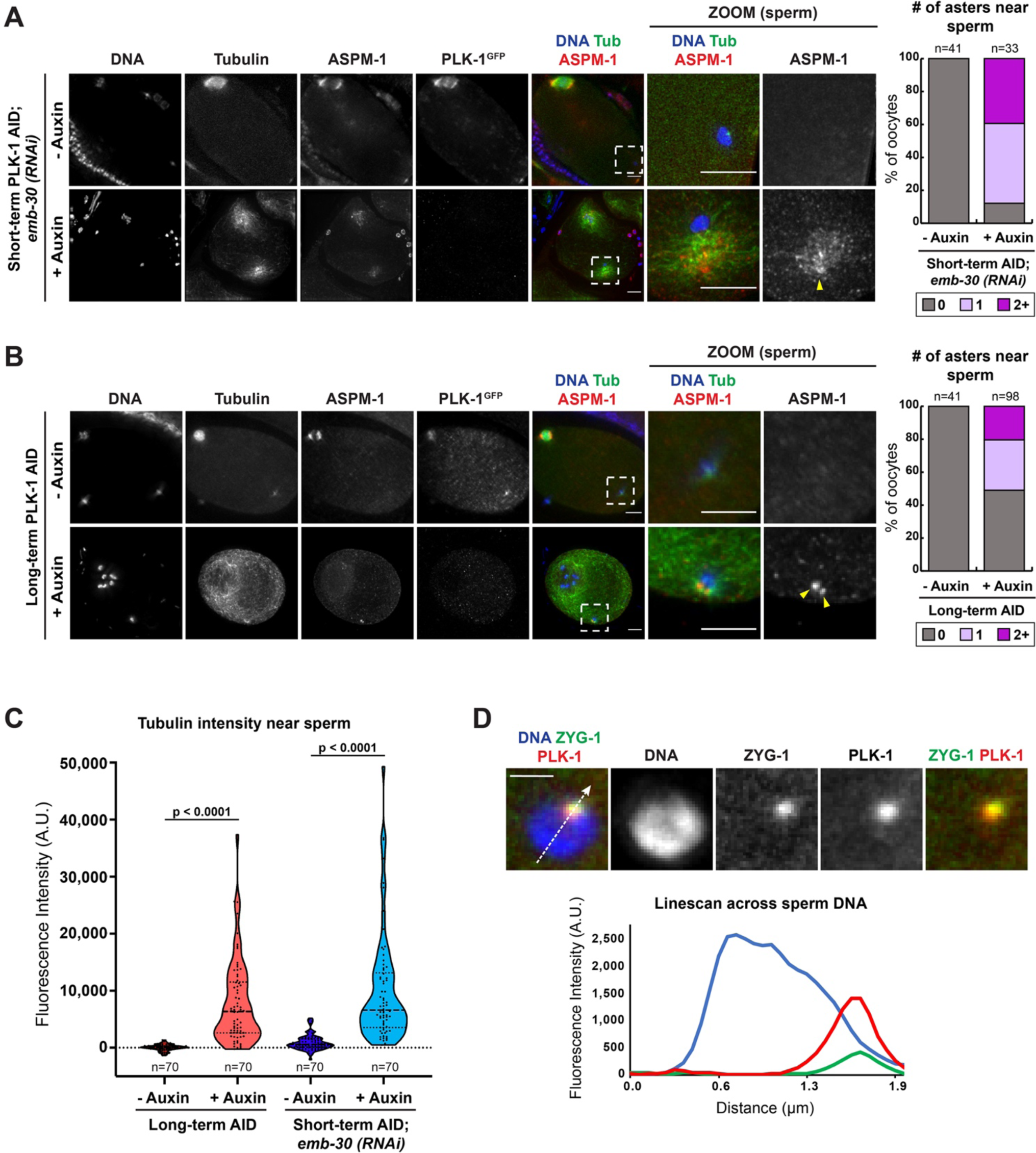
PLK-1 localizes to the sperm centriole and inhibits the formation of ectopic microtubule asters. A) Control and short-term auxin treated PLK-1 AID *emb-30(RNAi)* metaphase-arrested oocytes were stained for DNA (blue), tubulin (green), ASPM-1 (red) and PLK-1 (using a GFP antibody; not shown in merge) and imaged at 40X (columns 1-5) and 100X magnification (zooms). Quantification of ASPM-1 positive microtubule asters adjacent to sperm DNA shows an increase in ASPM-1 around the sperm coincident with microtubule density (arrowhead) in auxin treated oocytes. Scale bars = 5 μm. B) Control and long-term auxin treated PLK-1 AID oocytes were stained for DNA (blue), tubulin (green), ASPM-1 (red) and PLK-1 (using a GFP antibody; not shown in merge) and imaged at 40X (columns 1-5) and 100X magnification (zooms). Quantification of ASPM-1 positive microtubule asters adjacent to sperm DNA (arrowheads) illustrates higher frequency of ectopic asters in long-term auxin treated oocytes compared to controls. Scale bars = 5 μm. C) Fluorescence intensity of tubulin around sperm DNA following long-term PLK-1 depletion (red) and short-term depletion from *emb-30(RNAi)* metaphase-arrested oocytes (blue). Higher fluorescence following auxin treatment indicates the presence of increased microtubule density around sperm DNA in PLK-1 depleted oocytes. Statistical significance was determined using the Mann Whitney U test. **** = P<0.0001. D) Immunofluorescence of PLK-1 (red) and ZYG-1 (green) near sperm DNA (blue) reveals localization of PLK-1 to the sperm centriole. Line scans (dashed white line) of PLK-1 and ZYG-1 fluorescence intensities on single slice projections show co-incident peaks in n=21 samples. Scale bar = 2.5 μm.

To determine if PLK-1 normally inhibits centrosome maturation in oocytes, we assessed whether PCM components were recruited to the vicinity of the sperm DNA after PLK-1 AID. We first investigated the localization of SPD-5, a central structural scaffold protein involved in recruiting downstream PCM components (Hamill, Severson, Carter, & Bowerman, 2002). Although SPD-5 normally does not localize to sperm centrioles until the completion of the meiotic divisions (McNally et al., 2012), we observed distinct SPD-5 foci adjacent to the sperm DNA following both short-term and long-term PLK-1 depletion (**Figure 6A, 6B**). We quantified the number of SPD-5 positive foci per oocyte and found that these foci are present in over 80% of oocytes in both AID conditions (**Figure 6A, 6B**). We speculate that instances of two foci represent SPD-5 localizing to the sperm-derived mother and daughter centrioles, and that one focus represents cases where we cannot resolve the centrioles from each other.

**Figure 6:**
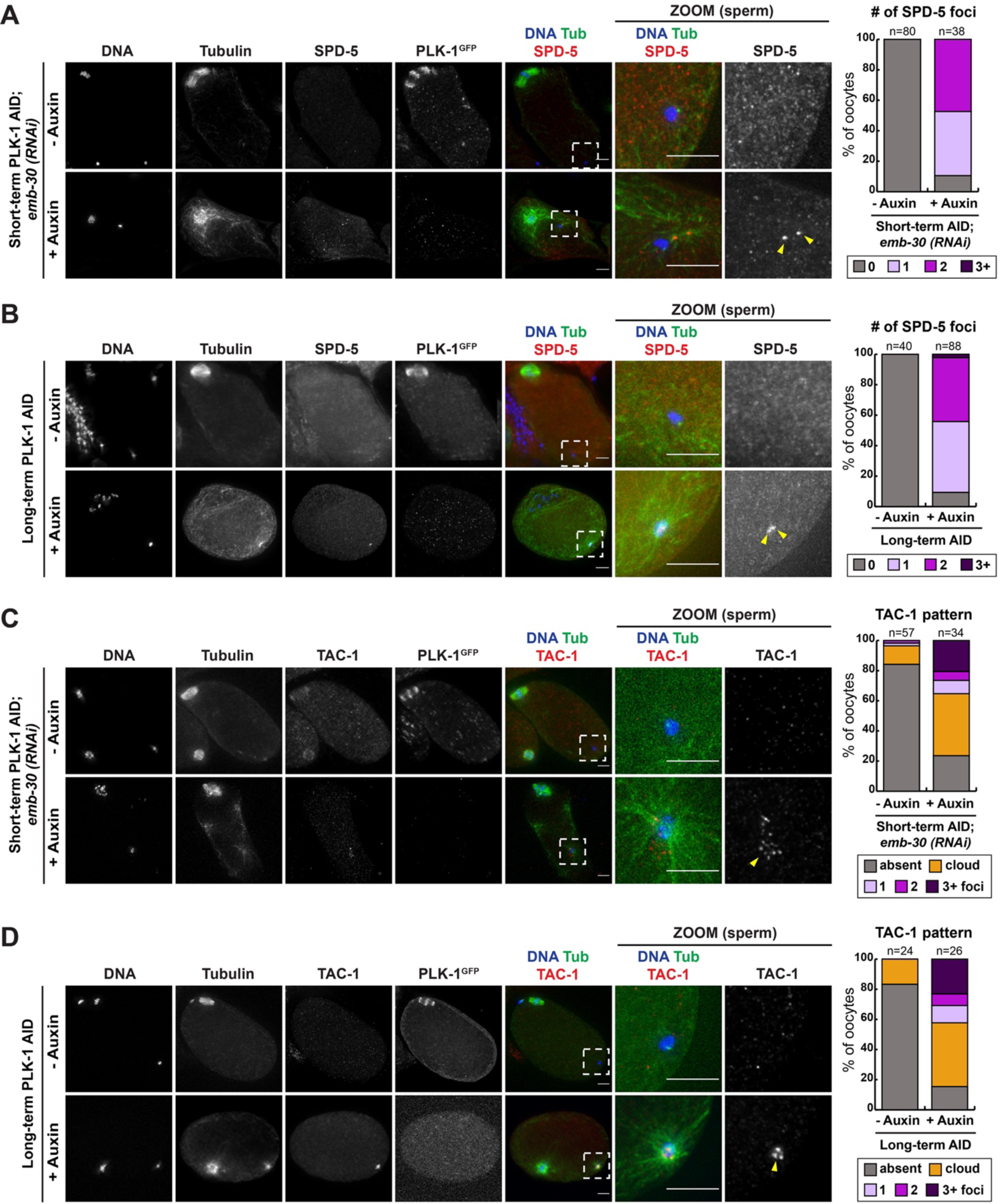
PLK-1 prevents premature recruitment of PCM components to the sperm-derived centrioles. A) Control and short-term auxin treated PLK-1 AID *emb-30(RNAi)* metaphase-arrested oocytes were stained for DNA (blue), tubulin (green), SPD-5 (red) and PLK-1 (using a GFP antibody; not shown in merge). All conditions in this figure were imaged at 40X (columns 1-5) and 100X magnification (zoomed images of the sperm DNA region). Quantification of number of SPD-5 puncta per cell indicates that oocytes treated with short-term auxin had increased SPD-5 localization near sperm DNA (arrowheads). B) Control and long-term auxin treated PLK-1 AID oocytes were stained for DNA (blue), tubulin (green), SPD-5 (red) and PLK-1 (using a GFP antibody; not shown in merge). Quantification shows an increased incidence of SPD-5 localization near sperm DNA after long-term PLK-1 AID (arrowheads). C) Control and short-term auxin treated PLK-1 AID *emb-30(RNAi)* metaphase-arrested oocytes were stained for DNA (blue), tubulin (green), TAC-1 (red) and PLK-1 (using a GFP antibody; not shown in merge). Categorization of TAC-1 localization pattern after short-term AID shows increased TAC-1 recruitment to sperm DNA after PLK-1 depletion. D) Control and long-term auxin treated PLK-1 AID oocytes were stained for DNA (blue), tubulin (green), TAC-1 (red) and PLK-1 (using a GFP antibody; not shown in merge). Corresponding quantification of the TAC-1 localization pattern indicates that long-term depletion of PLK-1 results in an increased localization of TAC-1 near sperm DNA. All scale bars = 5 μm.

We next assessed the localization of TAC-1, a PCM component associated with microtubule assembly in the *C. elegans* embryo (Le Bot et al., 2003). While TAC-1 does not typically concentrate near the sperm (McNally et al., 2012), in both short-term and long-term PLK-1 AID conditions we observed distinct foci of TAC-1 adjacent to the sperm DNA (**Figure 6C, 6D**). Both the TAC-1 and SPD-5 foci were often coincident with sperm-adjacent microtubule asters (**Figure 6A-D**). Taken together, these results demonstrate that depletion of PLK-1 leads to premature recruitment of PCM components to the sperm centrioles, indicative of premature centrosome maturation.

Notably, we did not observe SPD-5 or TAC-1 concentrated at the microtubule asters present in other regions of the oocyte (n=0/33 oocytes; n=1/16 oocytes, respectively) (**Figure S3**). These asters are therefore compositionally different from the sperm-adjacent asters, suggesting that they do not arise from PCM recruitment. Thus, PLK-1 appears to regulate microtubule assembly in the oocyte via multiple distinct mechanisms.

### PLK-1 depletion results in altered localization of KCA-1 around sperm DNA and shortened distances between the oocyte spindle and the sperm

Suppression of centrosome maturation during oocyte meiosis is essential to prevent premature sperm asters from interfering with the meiotic divisions (McNally et al., 2012).

Although the mechanisms suppressing centrosome maturation in *C. elegans* oocytes are poorly understood, a previous study found that depletion of either kinesin-1 or its cargo adaptor KCA-1 causes premature recruitment of PCM to the sperm centrioles (McNally et al., 2012). Therefore, the effects of PLK-1 depletion on the sperm centrioles could be related to misregulation of these proteins. Consistent with this hypothesis, we found that short-term and long-term depletion of PLK-1 resulted in altered localization of KCA-1 (**Figure 7A-7D**). In control conditions KCA-1 either forms a cloud around the sperm DNA, which has been proposed to directly inhibit PCM recruitment to the sperm centrioles, or does not display localization near the sperm, depending on stage of cell division (McNally et al., 2012) (**Figure 7A**, **7B**). Following short-term (**Figure 7A**, **7C**) and long-term (**Figure 7B**, **7D**) PLK-1 depletion, KCA-1 instead appeared as either 1 or 2 distinct puncta near the sperm in most oocytes. Thus, PLK-1 appears to regulate KCA-1, and presumably kinesin-1, localization in oocytes.

**Figure 7:**
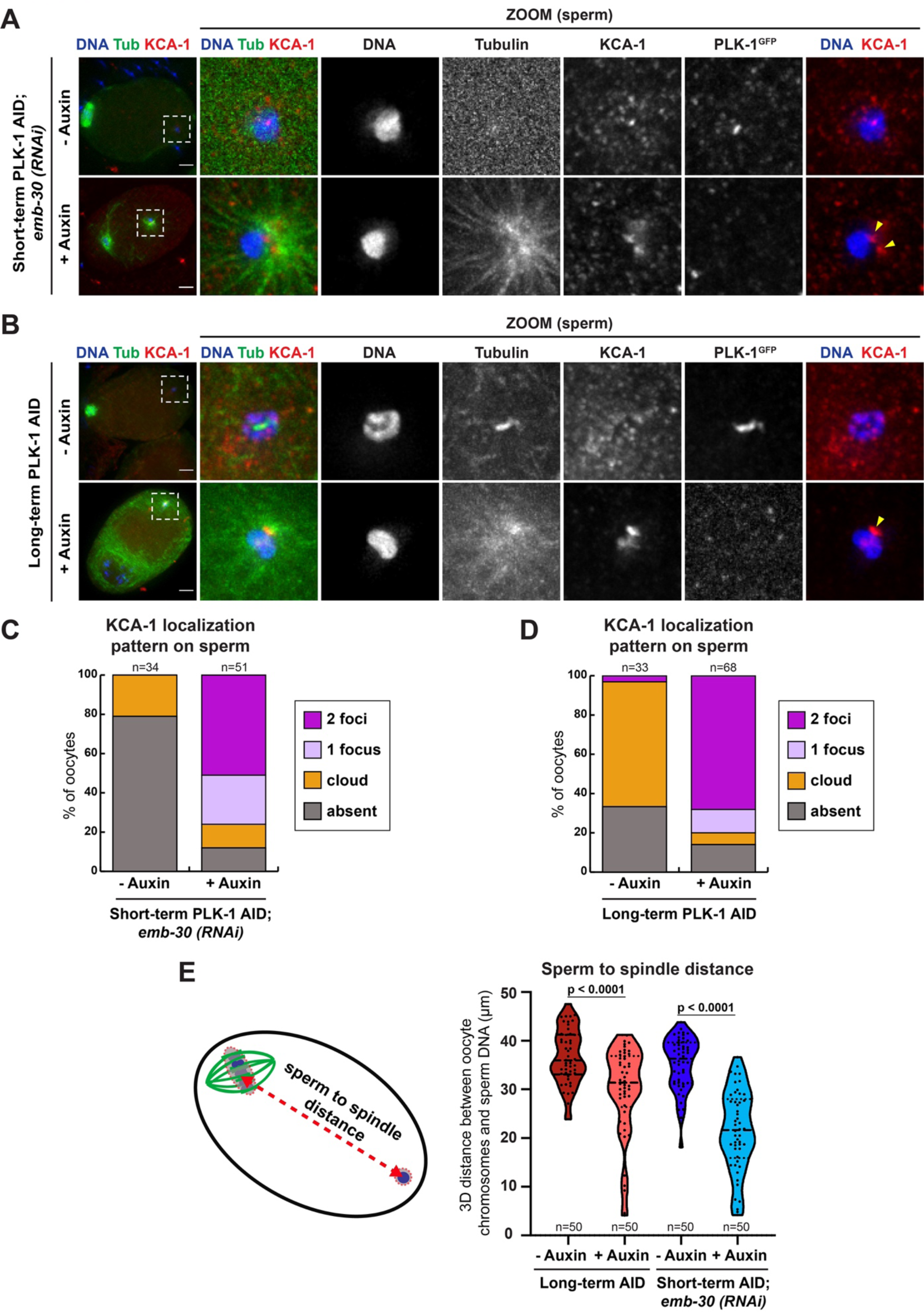
PLK-1 depletion results in altered KCA-1 localization around sperm DNA and a shortened distance between the oocyte chromosomes and sperm. A) Control and short-term auxin treated PLK-1 AID *emb-30(RNAi)* metaphase-arrested oocytes were stained for DNA (blue), tubulin (green), KCA-1 (red) and PLK-1 (using a GFP antibody; not shown in merge). All conditions in this figure were imaged at 40X (columns 1-5) and 100X magnification (zooms). Scale bars = 5 μm. B) Control and long-term auxin treated PLK-1 AID oocytes were stained for DNA (blue), tubulin (green), KCA-1 (red) and PLK-1 (using a GFP antibody; not shown in merge). Scale bars = 5 μm. C) Quantification of the experiment shown in A. KCA-1 typically forms a diffuse cloud around the sperm DNA or is absent, but after short-term auxin treatment KCA-1 re-localizes to 1 or 2 distinct puncta. D) Quantification of the experiment shown in B. Upon long-term PLK-1 depletion, KCA-1 re-localizes to distinct puncta near the sperm DNA. E) Schematic representation and 3D distance measurements between oocyte chromosomes and sperm DNA following long-term PLK-1 depletion (red) and short-term depletion from *emb-30(RNAi)* metaphase-arrested oocytes (blue). Distance was calculated by finding the center of the oocyte chromosome volume and the center of the sperm DNA volume and measuring the 3D distance between both points. Decreased distances between oocyte chromosomes and sperm DNA were observed following auxin treatment. Statistical significance was determined using the Mann Whitney U test. **** = P<0.0001.

To further explore a connection between PLK-1 and kinesin-1/KCA-1, we assessed the distance between the sperm DNA and the oocyte spindle in PLK-1 depleted oocytes. A previous study demonstrated that depletion of KCA-1 caused these structures to move closer together, sometimes resulting in capture of the meiotic spindle by the sperm aster (McNally et al., 2012). Similarly, we found that PLK-1 depletion shortened the average distance between the sperm DNA and the oocyte chromosomes in both long-term (29.8 µm) and short-term AID conditions (21.5 µm) compared to controls (36.6 µm and 35.2 µm, respectively; **Figure 7E**). We also found cases where the sperm aster was adjacent to the oocyte chromosomes, suggesting a role for PLK-1 in ensuring that the meiotic spindle and sperm DNA remain positionally separate within the oocyte. These findings support the interpretation that by preventing premature centrosome maturation, PLK-1 allows the meiotic divisions to be completed without interference.

## DISCUSSION

### PLK-1 plays multiple important roles during oocyte meiosis

Taken together, our work shows that PLK-1 fulfills a number of important roles during oocyte meiosis (**Figure 8**). PLK-1 dynamically localizes to various regions of the oocyte spindle, where it is required for both bipolar spindle assembly and stability. Moreover, we found that PLK-1 regulates microtubule polymerization in oocytes; this kinase prevents excess microtubule polymerization throughout the entire oocyte, and also localizes to the sperm-provided centrioles where it prevents premature centrosome maturation. Overall, these results demonstrate that PLK-1 promotes faithful meiosis in multiple distinct ways.

**Figure 8:**
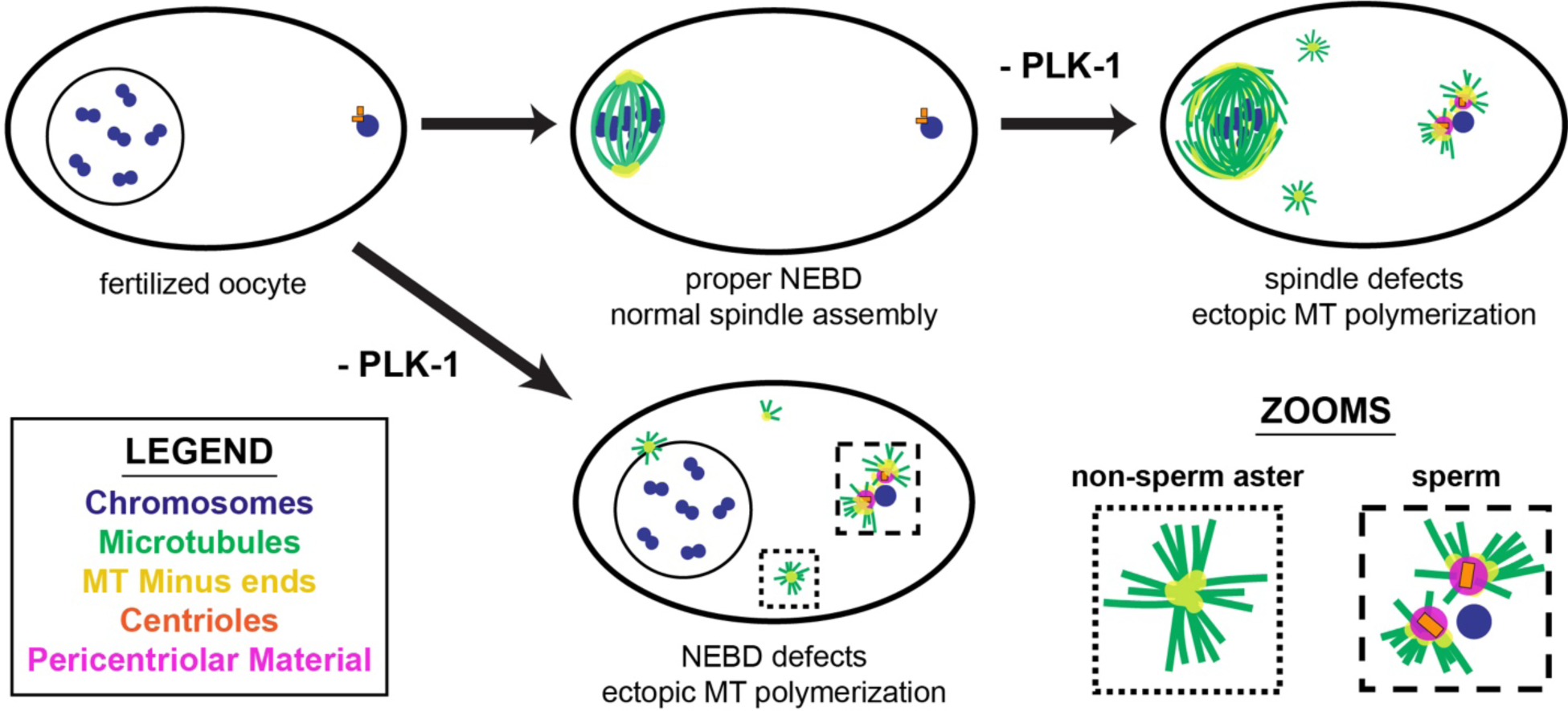
PLK-1 plays several distinct roles in meiosis, enabling nuclear envelope disassembly, bipolar spindle maintenance and inhibition of premature centrosome maturation. Model figure summarizing the effects of PLK-1 depletion on DNA (blue), microtubules (green), microtubule minus ends (yellow), centrioles (orange), and pericentriolar material (magenta). Upon long-term depletion of PLK-1, defective nuclear envelope breakdown occurs, preventing reliable bipolar spindle assembly. When PLK-1 is present at fertilization, the nuclear envelope breaks down (NEBD) and a bipolar spindle assembles. Depletion of PLK-1 after bipolar spindle formation results in disorganized spindles with microtubule (MT) minus ends distributed throughout the structure rather than concentrated at two distinct poles. In PLK-1 depleted oocytes ectopic microtubule polymerization occurs throughout the cell, including adjacent to the sperm DNA where PCM is prematurely recruited.

### PLK-1 promotes aMTOC-independent spindle assembly and regulates microtubule polymerization in oocytes

In mouse oocytes, aMTOCs comprised of PCM proteins nucleate microtubules and form asters that assist with spindle assembly; aMTOCs collect along the nuclear envelope and fragment into smaller structures upon NEBD that distribute to the two spindle poles (Luksza, Queguigner, Verlhac, & Brunet, 2013; Schuh & Ellenberg, 2007). It has been shown that PLK-1 is required for aMTOC fragmentation, thus facilitating proper spindle formation (Clift & Schuh, 2015; Little & Jordan, 2020; Solc et al., 2015). Now, we report that PLK-1 also is required for spindle assembly in *C. elegans* oocytes, a system that is not thought to rely on aMTOCs (Connolly et al., 2015; Wolff et al., 2016). Thus, PLK-1 contributes to acentrosomal spindle assembly in multiple ways.

Notably, although aMTOCs have not been previously observed in *C. elegans* oocytes, we observed microtubule asters throughout the oocyte after PLK-1 depletion. Thus, we considered the possibility that these asters might be transient aMTOCs that are normally difficult to image, but become large and/or stable enough to visualize upon PLK-1 depletion. To investigate this hypothesis, we assessed whether known aMTOC components localize to the ectopic asters. We found that SPD-5, whose homolog CDK5RAP2 localizes to aMTOCs in mouse oocytes (So et al., 2019), was not detectable in these ectopic asters. Similarly, the centrosome-associated protein TAC-1, whose homolog TACC3 localizes to a singular aMTOC during the early stages of spindle assembly in human oocytes (Wu et al., 2022), was also not present on the microtubule asters. Thus, if the asters observed upon PLK-1 depletion are stabilized aMTOCs, they are comprised of alternate proteins than the canonical proteins associated with aMTOCs in other systems.

While spindle defects were apparent soon after oocytes were dissected into auxin, visualizing ectopic microtubule aster formation required pre-treating worms prior to live imaging. This requirement could be for several reasons. One rationale is that full depletion of PLK-1 is necessary in order to have such a dramatic effect on the oocyte. Another possibility is that it takes time for PLK-1 depletion to influence downstream targets that in turn trigger excess microtubule polymerization. Microtubule growth has been linked to the amount of free tubulin in the cell (which should be the same in the oocyte regardless of PLK-1 AID) as well as the amount of ZYG-9 and TAC-1 (Srayko, Kaya, Stamford, & Hyman, 2005), which are associated with microtubule assembly in the *C. elegans* embryo (Bellanger & Gonczy, 2003). Since TAC-1 does not localize to the ectopic asters throughout the oocyte, we infer the same for ZYG-9 because these proteins co-localize in mitosis and meiosis (Cavin-Meza et al., 2022; Le Bot et al., 2003). Thus, we speculate that the asters throughout the oocyte may be due to altered activity of other microtubule regulatory factors. A plausible candidate for such a factor is the microtubule depolymerase KLP-7^MCAK^, as a previous study reported ectopic asters upon KLP-7 depletion (Gigant et al., 2017). However, because co-depletion of PLK-1 and KLP-7 had a more extreme phenotype than either single depletion alone, the increased microtubule polymerization seen upon PLK-1 depletion is unlikely to be solely due to KLP-7 misregulation. Other known PLK-1 targets include canonical microtubule nucleators γ-tubulin and TPXL-1^TPX2^ (Roostalu, Cade, & Surrey, 2015; Strome et al., 2001; Woodruff et al., 2017). These proteins could also be playing a role in this PLK-1 depletion phenotype, however, depletion of these proteins has not been reported to cause meiotic spindle defects in past studies (Bobinnec, Fukuda, & Nishida, 2000; Ozlu et al., 2005). Future work aimed at identifying PLK-1 targets that regulate microtubule polymerization in the oocyte may elucidate how ectopic aster formation is normally suppressed.

### The role of PLK-1 in suppressing centrosome maturation during oocyte meiosis

Another question raised by our findings is how PLK-1 suppresses the maturation of the sperm-derived centrioles during meiosis. This is especially puzzling since PLK-1 plays the opposite role during mitosis, where it facilitates the recruitment of SPD-5 and other PCM components to centrosomes (Lane & Nigg, 1996; Lee & Rhee, 2011). Conversely, in oocytes we find that PLK-1 localizes to the sperm-derived centrioles and appears to block PCM accumulation.

How PLK-1 performs these two opposite functions is a fascinating question for future study. Previous work demonstrated that kinesin-1 and its cargo adaptor KCA-1 prevent the accumulation of SPD-5 and other PCM components on the sperm centrioles prior to the completion of meiosis (McNally et al., 2012). Our finding that PLK-1 depletion results in altered KCA-1 localization raises the possibility that redistribution of these proteins triggers premature PCM recruitment. It has been proposed that kinesin-1/KCA-1 may act by forming a shell around the sperm DNA that blocks PCM assembly until the completion of the meiotic divisions (McNally et al., 2012). If this model is correct then disruption of this shell upon PLK-1 depletion would initiate premature centrosome maturation. In this scenario, PLK-1 could potentially act by phosphorylating kinesin-1 and/or KCA-1 to control their localization. Alternatively, PLK-1 could phosphorylate PCM components to block their accumulation during meiosis, or phosphorylate other regulatory factors. Further experiments examining known PLK-1 consensus motifs (Nakajima, Toyoshima-Morimoto, Taniguchi, & Nishida, 2003) and manipulating putative phosphorylation sites in candidate proteins may help elucidate the mechanism of how PLK-1 depletion is affecting premature centrosome maturation.

Another fascinating question relates to the mechanism by which SPD-5 accumulates at the sperm-derived centrioles in the absence of PLK-1. During centrosome maturation in *C. elegans* mitosis, SPD-5 oligomerizes and builds a scaffold for the recruitment of other PCM components (Ohta et al., 2021; Woodruff et al., 2015; Wueseke et al., 2016). Since PLK-1-mediated phosphorylation is thought to drive SPD-5 multimerization (Nakajo, Kano, Tsuyama, Haruta, & Sugimoto, 2022; Rios et al., 2024) how does SPD-5 accumulate at centrioles and drive the recruitment of other PCM components if PLK-1 is not present? One possibility is that SPD-5 accumulates without being phosphorylated under these conditions, which would suggest that phosphorylation is not always required for SPD-5 multimerization. Alternatively, it is possible that another kinase may be able to phosphorylate SPD-5 when PLK-1 is depleted. Future experiments distinguishing between these possibilities will enhance our understanding of SPD-5 multimerization and centrosome assembly.

Interestingly, another known function of PLK-1 is to promote disengagement of the centriole pair prior to mitosis, a process that is required for subsequent PCM recruitment (Tsou et al., 2009). Consistent with PLK-1’s alternate role in our system, our data suggests that during oocyte meiosis PLK-1 may prevent premature separation of the two centrioles. Typically, the mother and daughter centriole disengage after the completion of meiosis II (Cabral, Sans, Cowan, & Dammermann, 2013); however, we often observed 2 distinct foci of SPD-5 and KCA-1 adjacent to the sperm DNA in PLK-1-depleted oocytes. While centrioles were not labeled in these conditions, we reason that these 2 distinct foci mark the mother and daughter centrioles as they are prematurely separating and recruiting PCM. This contrasts with the single focus of ZYG-1/PLK-1 observed in untreated oocytes, which we infer marks the closely associated centriole pair prior to disengagement.

In human cells, Plk1 and separase function at different times to activate centriole disengagement during mitosis (Kim, Lee, & Rhee, 2015; Tsou et al., 2009). The *C. elegans* ortholog of separase, SEP-1, has also been shown to promote separation of the sperm-derived centrioles after the completion of female meiosis (Cabral et al., 2013). Interestingly, while *sep-1(RNAi)* results in defective centriole separation during mitosis, co-depletion of SEP-1 and KCA-1 via RNAi rescues this phenotype (Cabral et al., 2013), indicating a potential link between centriole separation and KCA-1. Given that we observed altered localization of KCA-1 following PLK-1 depletion as well as centriole disengagement, we postulate that premature centriole disengagement may be caused by KCA-1 mislocalization.

In summary, reported in this study are multiple novel roles for PLK-1 in oocyte meiosis that are distinctly different from the canonical roles of this kinase in mitosis. These alternate roles for PLK-1 during oocyte meiosis provide abundant opportunities for follow-up investigations on meiotic spindle dynamics, microtubule polymerization, and centrosome maturation.

## MATERIALS AND METHODS

### *C. elegans* strain generation and maintenance

Strains PHX3354 and PHX7793 were generated by SunyBiotech via CRISPR/Cas9 editing of the endogenous *plk-1* locus in the CA1199 (Zhang et al., 2015) and SMW44 background strains respectively. All strains were maintained at 15°C.

### Immunofluorescence and antibodies

Immunofluorescence was performed as described in Wolff et al. 2022. Briefly, adult worms were picked into a 10 μL drop of Meiosis Medium (0.5 mg/mL Inulin, 25 mM HEPES, and 20% FBS in Leibovitz’s L-15 Media (Gibco 11415-048)) (Laband, Lacroix, Edwards, Canman, & Dumont, 2018) on poly-L-lysine slides and dissected to remove oocytes. Slides were covered with a coverslip and slowly lowered into liquid nitrogen for 5–10 minutes, after which the coverslip was rapidly removed via razor blade and the slide was submerged in –20°C MeOH for 45 minutes. Samples were rehydrated in PBS, blocked in AbDil (PBS with 4% BSA, 0.1% Triton-X-100, 0.02% Na-Azide) overnight at 4°C, and then incubated in primary antibodies overnight at 4°C. The following day slides were washed three times with PBST (PBS with 0.1% Triton-X-100) and incubated with secondary antibodies for two hours at room temperature. After three PBST washes, samples were incubated with mouse anti-α-Tubulin-FITC (Sigma, 1:500) for two hours at room temperature and washed again. Samples were then incubated with Hoescht (1:1000 in PBST) for 15 minutes and washed twice with PBST. Finally, samples were mounted in 0.5% p-phenylenediamine, 20 mM Tris-Cl, pH 8.8, 90% glycerol, sealed with nail polish and stored at 4°C.

The primary antibodies used in this study were mouse-α-Tubulin-FITC (1:500, DM1α, Sigma), mouse-α-GFP (1:250, 3E6, Invitrogen), rabbit-α-ASPM-1 (1:5000, gift from Arshad Desai), rabbit-α-TAC-1 (1:50, (Cavin-Meza et al., 2022)), rabbit-α-SPD-5 (1:1500, gift from Bruce Bowerman) and rabbit-α-KCA-1 (1:200, gift from Frank McNally). ChromoTek Spot-Label 568 (1:800, Proteintech) was used to detect ZYG-1. Alexa Fluor rabbit and mouse secondary antibodies (Invitrogen) were all diluted 1:500 in PBST.

### RNAi feeding

RNAi was performed as described in Wolff et al., 2022. Briefly, individual RNAi cultures were grown by picking individual clones from a RNAi library (Fraser et al., 2000; Kamath et al., 2003) at 37°C in LB supplemented with 100μg/mL ampicillin. Cultures were grown overnight, centrifuged, and plated on nematode growth medium (NGM) plates containing 100 μg/mL ampicillin and 1 mM IPTG. Plates were dried overnight at room temperature in the dark. Worms were synchronized in preparation for experimentation by bleaching gravid adults, collecting resulting embryos and incubating on plates lacking food overnight. The following day, hatched L1s were transferred to RNAi plates and grown to adulthood at 15°C for 6 days.

### Auxin treatment

Multiple modes of auxin treatment were utilized in this study, as summarized in **Figure 1A** and briefly described here. Further details are available in Divekar et al., 2021.

#### Long-term AID

Long-term auxin was administered by incubating worms on NGM plates containing auxin for 18 hours. Plates were prepared as normal NGM plates except for the addition of auxin dissolved in 100% EtOH for a final concentration of 1 mM auxin. Plates were stored in the dark at 4°C for no longer than 2 months. For experimentation, synchronized worms (described earlier) were grown to L4s on standard NGM plates. Worms were then transferred onto 1 mM auxin containing plates and incubated at 15°C for 18 hours prior to dissection and immunofluorescence.

#### Short-term AID

Short-term auxin treatment was performed by soaking intact worms in Meiosis Media containing 5 mM auxin. This solution was prepared by diluting a stock of 400 mM auxin dissolved in 100% EtOH that was protected from light. For experimentation, adult worms were picked into 10 μL of 5 mM auxin solution and incubated for 40 minutes in a humidity chamber to prevent evaporation of the solution. For vehicle control treatment, adults were picked into 10 μL of Meiosis Media with the equivalent volume of 100% ethanol and incubated identically. Following 40 minutes of incubation, worms were dissected (∼5 minutes) and then progressed through the standard immunofluorescence protocol as described above for a total of ∼45 minutes of auxin treatment. *Acute AID:*

Acute auxin treatment was performed by picking 10-15 worms into 10 µL of Meiosis Media containing 500 µM auxin. This solution was prepared using a stock of 400 mM auxin dissolved in 100% EtOH. The control solution consisted of an equivalent volume of 100% ethanol diluted in Meiosis Media. Adult worms were picked into a 10 μL drop of auxin or control solution and immediately dissected and mounted for *ex utero* live imaging as described below.

#### Acute AID + pre-treatment

Worms were set up identically to the acute AID protocol, except for one additional incubation step. After worms were picked into 10 μL of solution, the entire slide was incubated in a humidity chamber for 30 minutes at room temperature to prevent evaporation of the liquid. This incubation time allowed for live imaging visualization of the oocyte after a more complete PLK-1 depletion as compared to acute AID. After this pre-treatment step, worms were immediately dissected and mounted for *ex utero* live imaging.

### *Ex utero* Live Imaging

Worms were taken from desired experimental plates and dissected into 10 µL of auxin or control solution placed in the center of a live imaging apparatus (Divekar et al., 2021; Laband et al., 2018). A Vaseline ring was made using a syringe to contain the sample, and a 18x18 mm coverslip was laid on top. The slide was inverted and immediately imaged.

### Microscopy

Fixed imaging was performed on a DeltaVision Core deconvolution microscope using either a 40X (NA = 1.3) or 100X objective lens (NA = 1.4) (Applied Precision). Z-stacks were obtained at 0.2 μm increments, deconvolved using SoftWoRx (Applied Precision), and processed using ImageJ. Images are shown as maximum intensity projections unless otherwise indicated. This microscope is housed in the Northwestern University Biological Imaging Facility supported by the NU Office for Research.

Live imaging was performed using a Nikon SoRa spinning disk confocal microscope with an oil-immersion 60x (1.42 NA) objective lens. Images were acquired using a Yokogawa CSU-W1 dual-disk spinning disk unit with a 50 μm pinhole, and a Hamamatsu ORCA-Fusion Digital CMOS Camera. The microscope was controlled by the Nikon SoRa imaging software NIS-Elements AR. Fourteen z-stacks at 0.5 μm increments were taken every 30 seconds at room temperature. Videos were processed using ImageJ to create maximum intensity projections. The Nikon SoRa microscope is housed in the Northwestern University Biological Imaging Facility supported by the NU Office for Research.

### Data analysis

When quantifying our results, similar phenotypes were observed in MI and MII. Therefore, all quantifications of phenotypic categories represent pooled results from MI and MII oocytes (**Figures 1D, 2B, 3C, 5A-B, 6A-D, 7C-D**).

#### Ectopic aster intensity (Figure 4C)

Sum projections were made of all z slices in the tubulin channel at t=12 minutes using ImageJ. Mean gray values ranged from 0 to 255, so a threshold intensity value was set at 64 (corresponds to dark blue in **Figure 4B**). The number of asters within the oocyte that exceeded this threshold value were counted. Additionally, the fluorescence intensity value of any area within the oocyte above the threshold was measured using a ROI set to reflect the average aster size (approximately 4 μm x 4 μm). The same ROI was used to analyze all samples.

#### Tubulin intensities near sperm (Figure 5C)

Using 40X images of the entire oocyte, sum projections of tubulin intensity were made in ImageJ from 1.4 μm above and below the center of the sperm DNA (2.8 μm stack total, 14 z-slices). The fluorescence intensity was measured surrounding the sperm DNA using a ROI of approximately 4 μm x 4 μm. The same ROI was used to measure three other areas within the oocyte cytoplasm (away from the meiotic spindle and sperm DNA) for background subtraction of cortical tubulin intensity. The average tubulin intensity of the background was subtracted from the tubulin intensity surrounding the sperm DNA, and the resulting values were graphed.

#### PLK-1 and ZYG-1 co-localization (Figure 5D)

Fluorescence intensities for DNA, PLK-1, and ZYG-1 were measured using ImageJ on single-slice projections using the same 2 μM long line, drawn from one end of the sperm DNA to just past the center of the sperm centriole, as determined by ZYG-1 staining. Directionality of the line is indicated by the white arrow.

#### Sperm distance measurements (Figure 7F)

Using 40X images of the entire cell, oocyte chromosomes and sperm DNA were rendered into 3D surfaces using the ‘Surfaces’ tool in the 3D Imaging Software Imaris (Bitplane). For each cell analyzed, the center of both the oocyte chromosomes and sperm DNA was determined using the DAPI channel, and the distance between them was measured in 3D, as depicted in **Figure 7E**.

### Statistical Methodology

All statistics were performed using GraphPad Prism 10. Statistical methodology, significance values and sample sizes are indicated in corresponding figures. The Shapiro-Wilk Normality was used to test for normality of data distribution. Normally distributed data was analyzed using the two-tailed t-test, while non-normally distributed data was analyzed using the Mann Whitney U nonparametric test.

## ACKNOWLEDGMENTS

We would like to thank members of the Wignall lab for support, and Gabriel Cavin-Meza, Emily Czajkowski, Hannah Horton, Elizabeth Lopez, Ilka Lorenzo, and Jordy Martinez for critical reading of the manuscript. We are also grateful to Bruce Bowerman, Frank McNally, and Kevin O’Connell for reagents. This work was supported by NIH R01GM124354 and NIH R01GM141386 (to SMW) and by the NIH Reproductive Science, Medicine, and Technology training grant T32HD094699 (to JGN). Microscopy was performed at the Biological Imaging Facility at Northwestern University, supported by the NU Office of Research and the Department of Molecular Biosciences.

---

**Figure S1:**
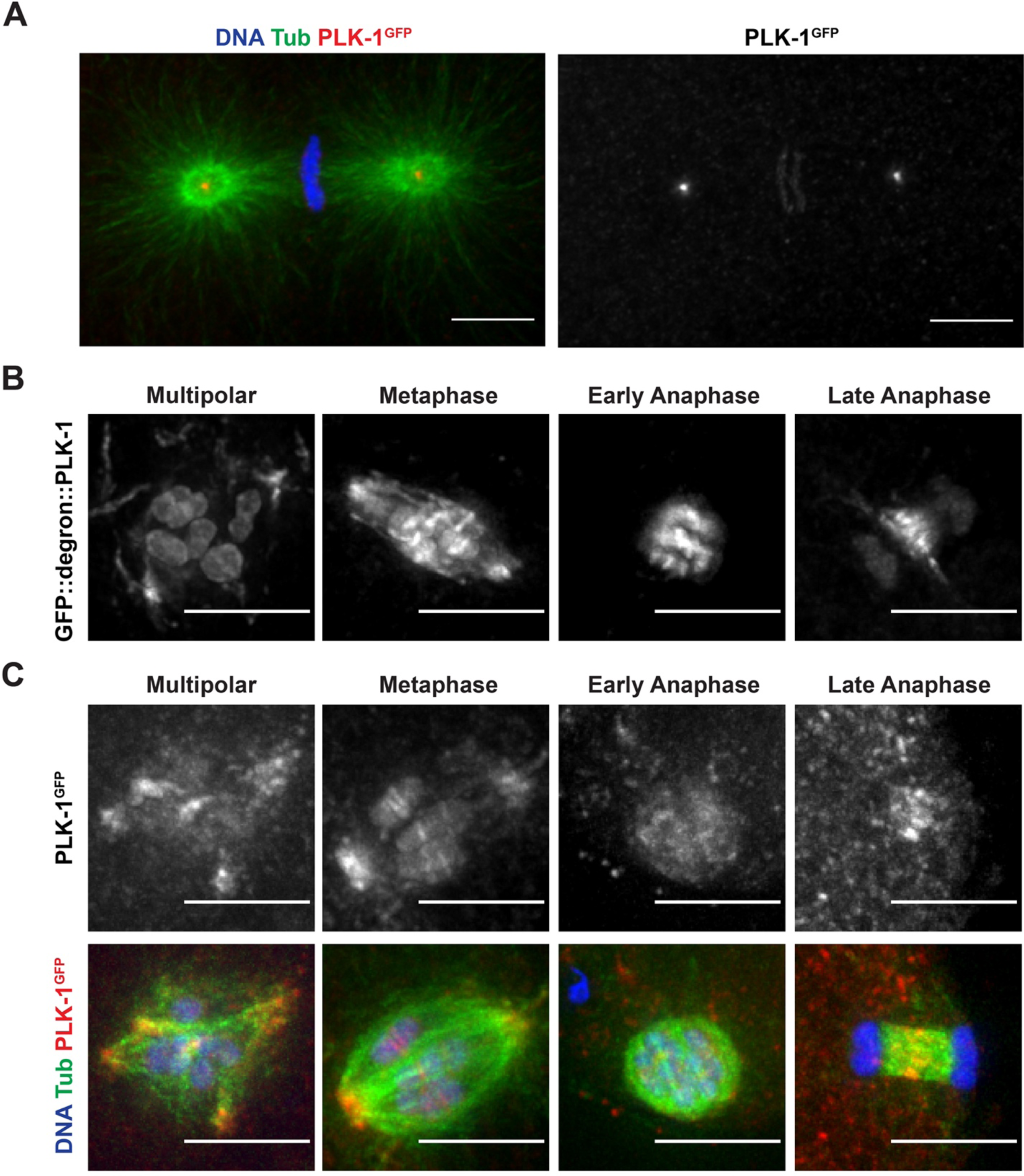
Localization of PLK-1 on the meiotic spindle during various stages of oocyte meiosis using an AID strain is consistent with existing literature. A) Immunofluorescence of an embryo from the PLK-1 AID strain using a GFP antibody shows that PLK-1 localizes to the centrosomes and kinetochores during mitosis. Shown are DNA (blue), tubulin (green) and PLK-1 (red). B) Endogenous localization of GFP in ethanol fixed whole worms expressing GFP::degron::PLK-1 shows that PLK-1 localizes to the spindle poles, ring complex, DNA, kinetochore cups and kinetochore filaments during the multipolar and metaphase stages. During early anaphase, PLK-1 is present on the DNA and ring complex, and during late anaphase PLK-1 is on the spindle midzone and diffusely on the DNA. C) Immunofluorescence on untreated, dissected oocytes shows that PLK-1 localizes to the spindle poles, ring complex, DNA and spindle midzone during various stages of oocyte meiosis. Shown are DNA (blue), tubulin (green) and PLK-1 (red, stained with a GFP antibody). All scale bars = 5 μm.

**Figure S2:**
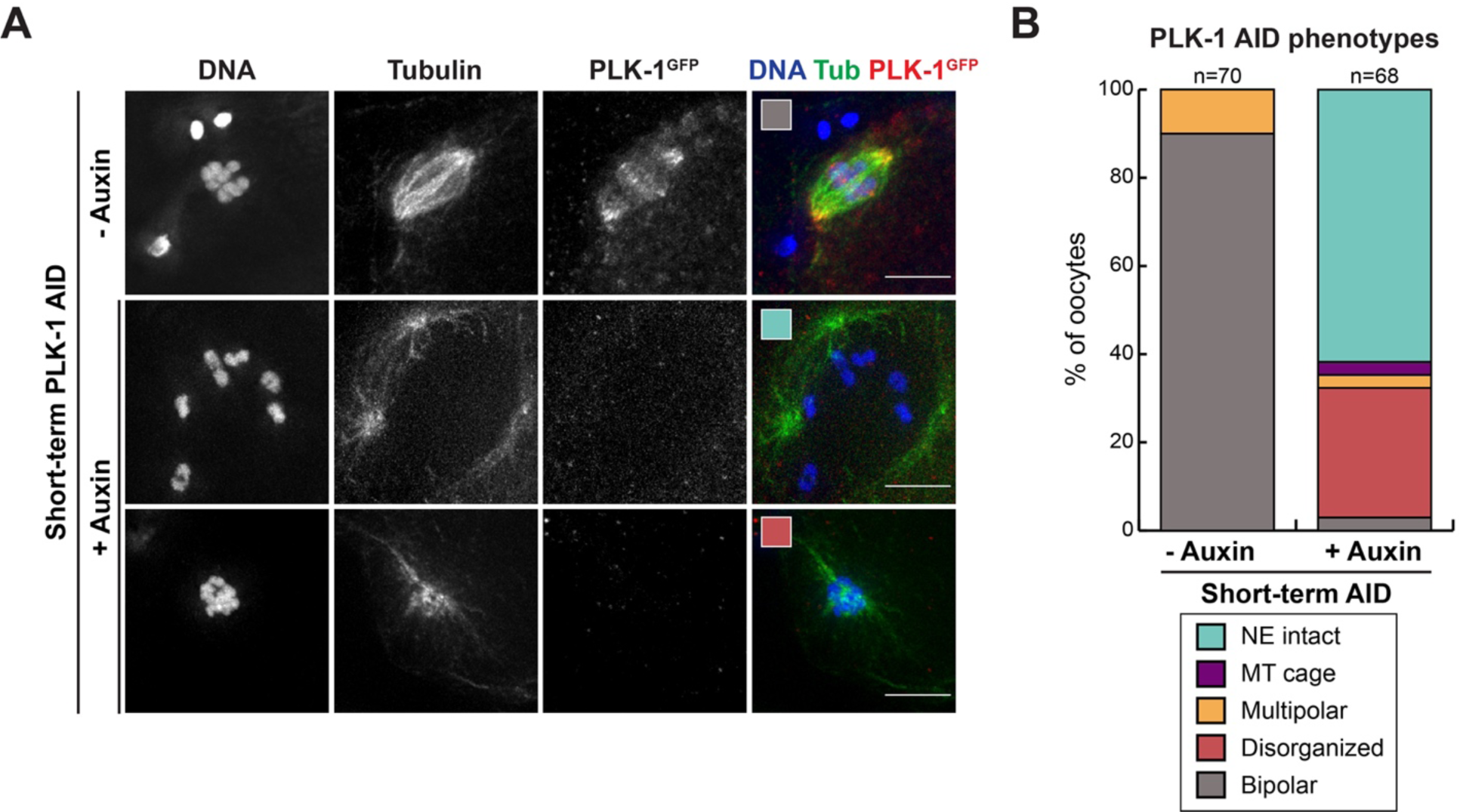
Short-term PLK-1 depleted oocytes fail to assemble and maintain bipolar spindles. A) Control and short-term auxin treated PLK-1 AID unarrested oocytes were stained for DNA (blue), tubulin (green) and PLK-1 (using a GFP antibody; red). Representative images of the major phenotypic categories quantified in B are indicated with colored boxes. Scale bars = 5 μm. B) Quantification of the experiment shown in A. Control spindles were largely bipolar, while short-term auxin treated oocytes were either disorganized or had an intact nuclear envelope, consistent with long-term AID.

**Figure S3:**
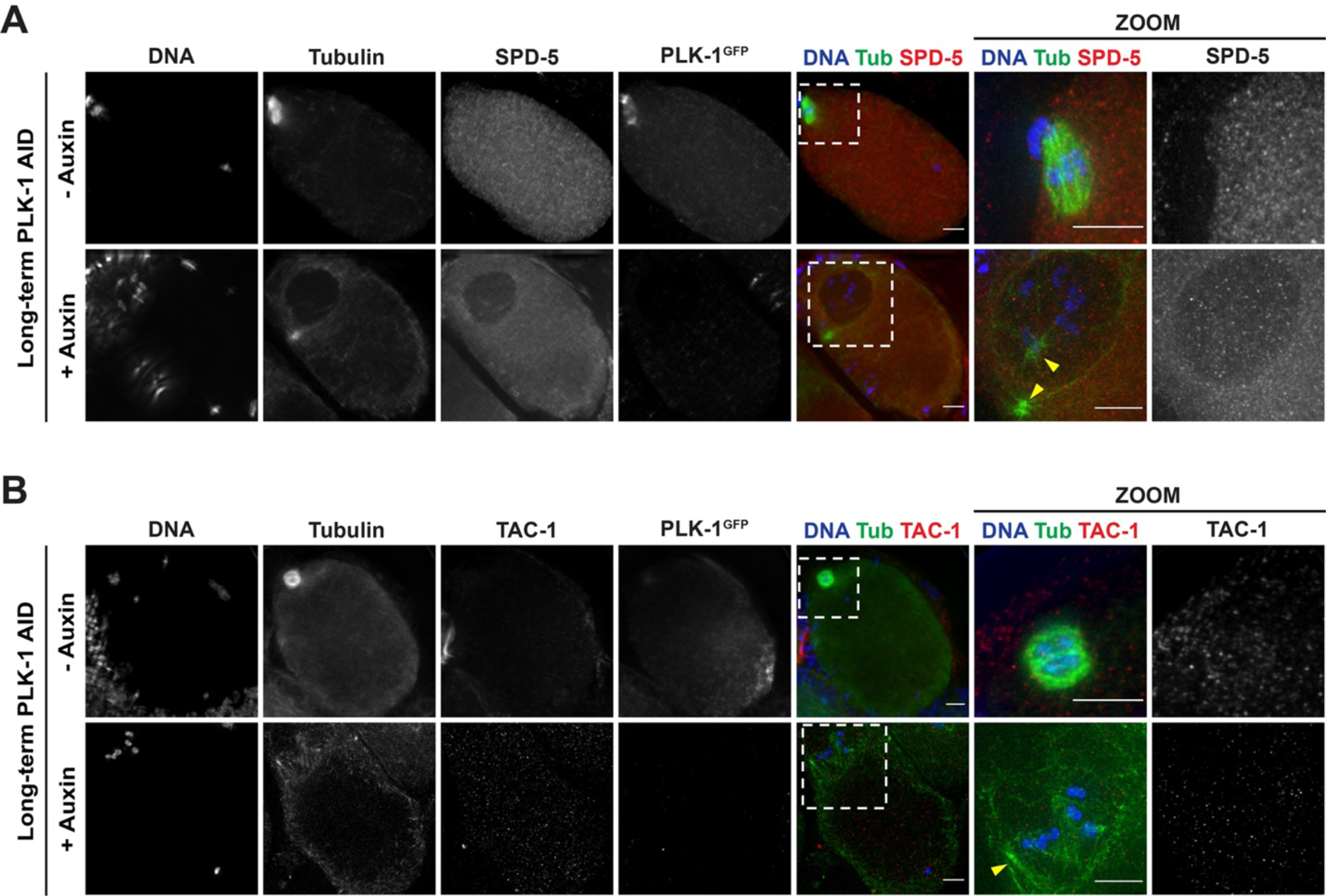
PCM components SPD-5 and TAC-1 are not present on oocyte chromosome-adjacent asters. Control and long-term auxin treated PLK-1 AID oocytes were stained for DNA (blue), tubulin (green), SPD-5 (red) and PLK-1 (using a GFP antibody; not shown in merge) (A) or DNA (blue), tubulin (green), TAC-1 (red) and PLK-1 (using a GFP antibody; not shown in merge) (B) and imaged at 40X (columns 1-5) and 100X magnification (zooms). SPD-5 localized to 0/33 asters (arrowheads) observed near the oocyte chromosomes in long-term auxin treated oocytes, while TAC-1 localized to 1/16 asters. Scale bars = 5 μm.

## SUPPLEMENTAL VIDEO LEGENDS

**Video 1: Metaphase-arrested oocyte spindles in the PLK-1 AID strain maintain bipolarity in the absence of auxin**

Live imaging of an *emb-30(RNAi)* metaphase-arrested oocyte spindle; corresponds to Figure 2C (top row). Shown are GFP::tubulin and GFP::PLK-1 (green), and mCherry::histone (magenta). Oocytes were dissected into Meiosis Media containing vehicle and immediately filmed. The spindle maintains bipolarity; chromosomes oscillate between either pole and the midspindle retains its integrity. This phenotype was consistent in all videos (n=7). Scale bar = 5µm.

**Video 2: Metaphase-arrested oocyte spindles in the PLK-1 AID strain elongate and lose midspindle integrity upon acute auxin treatment**

Live imaging of an *emb-30(RNAi)* metaphase-arrested oocyte spindle treated with auxin; corresponds to Figure 2C (bottom row). Shown are GFP::tubulin and GFP::PLK-1 (green), and mCherry::histone (magenta). Oocytes were dissected into auxin-containing Meiosis Media and immediately filmed. The spindle poles immediately move apart, the chromosomes lose alignment, and the midspindle splays resulting in a disorganized spindle. This phenotype was consistent in all videos (n=6). Scale bar = 5µm.

**Video 3: Unarrested oocyte spindles maintain bipolarity and undergo anaphase in the absence of auxin**

Live imaging of a control unarrested oocyte spindle; corresponds to Figure 2D (top row). Shown are GFP::tubulin and GFP::PLK-1 (green), and mCherry::histone (magenta). Oocytes were dissected into Meiosis Media containing vehicle and immediately filmed. The spindle maintains bipolarity in metaphase, shortens, rotates towards the cortex and then elongates to undergo anaphase. This phenotype was consistent in all videos (n=8). Scale bar = 5µm.

**Video 4: Unarrested oocytes exhibit spindle defects following acute PLK-1 AID**

Live imaging of an unarrested auxin treated oocyte spindle; corresponds to Figure 2D (bottom row). Shown are GFP::tubulin and GFP::PLK-1 (green), and mCherry::histone (magenta).Oocytes were dissected into auxin-containing Meiosis Media and immediately filmed. Upon auxin treatment the spindle elongates, chromosomes become misaligned and midspindle loses integrity. Spindle defects were observed in all videos (n=7). Scale bar = 5µm.

**Video 5: Pre-treatment of oocytes with vehicle does not yield excess tubulin densities**

Live imaging of a vehicle treated oocyte; corresponds to Figure 4A (top row). Shown are GFP::tubulin and GFP::PLK-1 (green), and mCherry::histone (magenta). Worms were soaked in vehicle-containing Meiosis Media for 30 minutes before oocytes were dissected and filmed. Control oocytes were able to complete meiosis I and successfully segregate chromosomes without spindle defects or excess tubulin density in the cell. This phenotype was consistent in all videos (n=7). Scale bar = 5µm.

**Video 6: Pre-treatment of oocytes with auxin results in ectopic microtubule polymerization**

Live imaging of an auxin treated oocyte; corresponds to Figure 4A (bottom row). Shown are GFP::tubulin and GFP::PLK-1 (green), and mCherry::histone (magenta). To achieve full PLK-1 depletion throughout the cell, worms were soaked in auxin-containing Meiosis Media for 30 minutes before oocytes were dissected and filmed. PLK-1 depleted oocytes formed tubulin-rich asters throughout the cell (n=7). Scale bar = 5µm.

**Video 7: *klp-7(RNAi)* oocytes exhibit some microtubule asters when PLK-1 is present**

Live imaging of tubulin density in a *klp-7(RNAi)* oocyte without auxin; corresponds to Figure 4B, middle row. Video is pseudocolored to show mean gray values of the GFP channel

(GFP::tubulin and GFP::PLK-1 signals). Oocytes were dissected into Meiosis Media containing vehicle and filmed. Control *klp-7(RNAi)* oocytes form some weak ectopic microtubule asters throughout the cell (n=8). Scale bar = 5µm.

**Video 8: No significant tubulin asters are present throughout the oocyte immediately following acute PLK-1 AID**

Live imaging of tubulin density in an auxin treated oocyte; corresponds to Figure 4B, top row. Pseudocolored video shows mean gray values of the GFP channel (GFP::tubulin and GFP::PLK-1 signals). Oocytes were dissected into Meiosis Media containing auxin and filmed. Microtubule asters were not observed immediately after acute AID since these oocytes were not pre-treated with auxin (n=7, same as Video 4 but zoomed out to view the entire oocyte). Scale bar = 5µm.

**Video 9: *klp-7(RNAi)* oocytes form many ectopic microtubule asters following acute PLK-1 AID**

Live imaging of tubulin density in a *klp-7(RNAi)* oocyte treated with auxin; corresponds to Figure 4B, bottom row. Pseudocolored video shows mean gray values of the GFP channel (GFP::tubulin and GFP::PLK-1 signals). Oocytes were dissected into auxin-containing Meiosis Media and filmed. Following auxin treatment, oocytes had increased tubulin density and multiple microtubule asters formed throughout the oocyte (n=8). Scale bar = 5µm.

